# Genome-wide Association study of facial morphology identifies novel genetic loci in Han Chinese

**DOI:** 10.1101/691584

**Authors:** Yin Huang, Dan Li, Lu Qiao, Yu Liu, Qianqian Peng, Sijie Wu, Manfei Zhang, Yajun Yang, Jingze Tan, Shuhua Xu, Li Jin, Sijia Wang, Kun Tang, Stefan Grünewald

## Abstract

Human face is a heritable surface with many complex sensory organs. In recent years, many genetic loci associated with facial features have been reported in different populations, yet there is a lack for the Han Chinese population. We report a genome-wide association analysis of 3D normal human faces in 2659 Han Chinese with two groups of phenotypes, the partial and whole face phenotypes and the distance and angle phenotypes. We found significant signals in five genomic regions with traits related to nose or eyes, including rs970797 in 2q31.1 near HOXD1 and MTX2, rs16897517 in 8q22.2 at intron of VPS13B, rs9995821 in 4q31.3 near DCHS2 and SFRP2, rs12636297 in 3q23 near PISRT1, and rs12948076 in 17q24.3 near SOX9 and CASC17. We visualized changes in facial morphology by comparing the volume of local areas and observed that these nose-related loci were associated with different features of the nose, including nose prominence, nasion height, and nostril shape, suggesting that the nose underlies precise genetic regulation. These results provide a more comprehensive understanding of the relationship between genetic loci and human facial morphology.

**Author Summary:** Human face as a combination of delicate sensory organs has a strong genetic component, as evidenced by the identical appearance in twins and shared facial features in close relatives. Although facial genetics have been studied in different populations, our knowledge between genetic markers with facial features is still limited. In this paper, we found genetic variants associated with nose and eyes through a large-scale high-resolution 3D facial genetic study on the Han Chinese population. We observed that these nose-related loci were associated with different features of the nose, including nose prominence, nasion height, and nostril shape, which suggests the nose underlies precise genetic regulation. Intriguingly, we noted that genes (*DCHS2* and *SFRP2*) related to one of these loci are differentially expressed in human and chimp cranial neural crest cells, which plays a crucial role in the early formation of facial morphology. The ongoing genetic studies of facial morphology will improve our understanding of human craniofacial development, and provide potential evolution evidence of human facial features.

## Introduction

The human face encompasses complex sensory organs and shows remarkable variation in facial traits. These complex and delicate traits undergo precise genetic regulation during facial development to produce our highly inherited, unique, and recognizable faces which play an essential role in communication and physical identification. Although the genetic basis of human facial morphology is still limited, many genetic variants associated with facial features have been reported in recent years.

In 2012, two studies (Liu *et al*., 2012; Paternoster *et al*., 2012) respectively found an association between a genetic variant in PAX3 with nose shape in European subjects, which was the start of GWAS of normal human facial morphology. Furthermore, Liu et al. identified four novel genetic variants close to the genes PRDM16, TP63, C5orf50, and COL17A1 affecting face morphology (Liu *et al*., 2012). More recently, a GWAS in Latin Americans reported five more genetic variants close to DCHS2, RUNX2, GLI3, PAX1, and EDAR related to nose shape (Adhikari *et al*., 2016). Another recent GWAS (Shaffer *et al*., 2016) identified seven additional genetic variants for face traits such as facial width and depth, and nose shape in European-derived cohort. Interestingly, this study observed the same association between soft-tissue nasal width and PAX1 identified in Latin Americans (Adhikari *et al*., 2016). Lee et al. identified two genetic variants of FREM1 and PARK2 associated with face shape in individuals of European ancestry (Lee *et al*., 2017). In 2018, two studies (Claes *et al*., 2018; Crouch *et al*., 2018) found a number of newly associated genetic loci by applying novel approaches to a European-derived cohort and replicated many loci in previous studies (Liu *et al*., 2012; Paternoster *et al*., 2012; Adhikari *et al*., 2016; Shaffer *et al*., 2016). In African subjects, SCHIP1 and PDE8A were identified to be associated with measures of human facial size (Cole *et al*., 2017). In Asians, a series of genome-wide association analyses of normal facial morphology on Uyghurs identified six significant loci (Qiao *et al*., 2018), and another study on Koreans identified five novel face morphology loci that were associated with facial frontal contour, nose shape, and eye shape (Cha *et al*., 2018).

So far, there were some GWAS studies carried out on normal human facial morphology in different populations, including Europeans (Liu *et al*., 2012; Paternoster *et al*., 2012; Shaffer *et al*., 2016; Lee *et al*., 2017; Claes *et al*., 2018; Crouch *et al*., 2018), Latin Americans (Adhikari *et al*., 2016), Africans (Cole *et al*., 2016) and Asians (Cha *et al*., 2018; Qiao *et al*., 2018). However, Studies on genome-wide associations of 3D normal human facial morphology in Han Chinese are still lacking.

In this study, we conducted genome-wide association analysis on the discovery cohort and the replication cohort for 8 partial-and-whole-face phenotypes and 10 distance-and-angle phenotypes by taking gender, age, and BMI as covariates in 2659 unrelated participants of HAN Chinese from Taizhou, Jiangsu province. We identified five genomic regions significantly associated with different facial phenotypes, including nose-related traits and eye-related traits. However, the relationship between genetic variants and facial morphology is still fuzzy due to the limitation of the representation of facial features. The face phenotypes tend to be either too simple to represent facial differences, or too complex to be intuitively explained. Therefore, we visualized the changes of local facial volume according to the characteristics of high-density registered 3D facial surface. Interestingly, we observed that different regions of the nose received precise regulation of genetic variants.

## Results

### Sample and phenotyping features

The study included 2659 unrelated participants of HAN Chinese with basic demographic descriptions and physiological characteristics (such as gender, age and BMI), which were recruited from Taizhou, Jiangsu province on the east coast of China. These participants had a mean age of 56.7 with an average BMI of 24.4 (SF1 A,D), 36% of which were male and 64% of which were female. The sample was randomly divided into discovery cohort (1766 participants) and replication cohort (893 participants) by two to one. Both cohorts had similar age (SF1 B,C) and BMI distributions (SF1 E,F) and sex ratio.

High-resolution 3D facial image was collected for each participant and automatically registered into a tri-mesh with 32251 points by an in-house landmark recognition method. Referred to the previous articles of our group (Qiao *et al*., 2018), two different schemes were used to characterize the facial morphologic phenotypes, including the partial and whole face phenotypes and the distance and angle phenotypes.

Briefly, the first scheme (the phenotypes of partial and whole face (PWF)) was extracted from high-density grid meshes with 32251 vertices. We roughly divided the face into different organic regions, including eyes, nose, cheeks, mouth and chin. Then a set of principle components explaining 97% of the total variance were extracted by PCA analysis as quantitative features representing the morphological information in the region (F1 A, SF2). For each face, two nose regions were extracted, one with only the nose (F1 B), and the other with the nose and adjacent regions (F1 C).

**Fig. 1.**
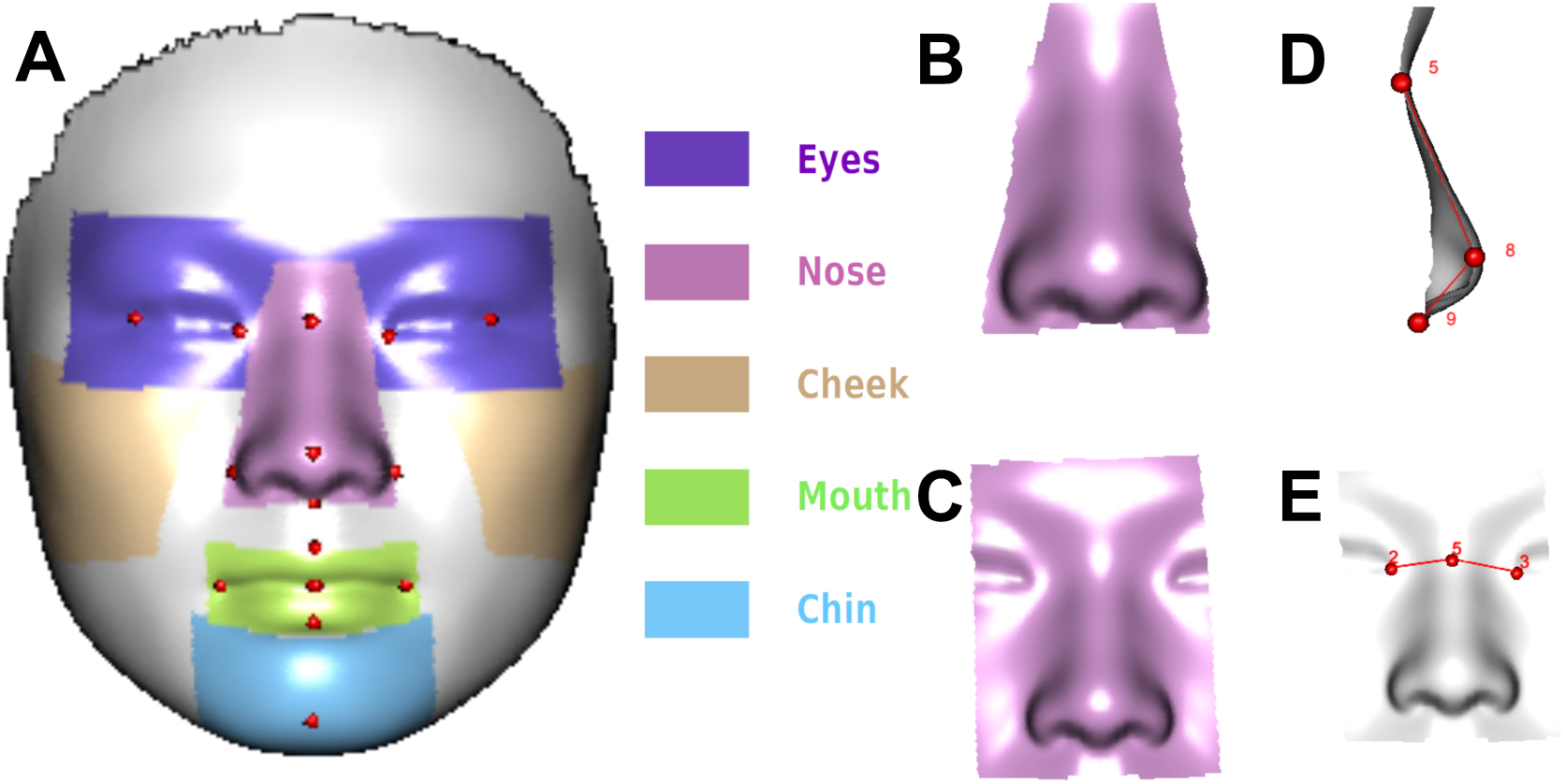
Two different schemes of the facial morphologic phenotypes. (A) The reference face with 15 landmarks is divided into eyes, nose, left cheek, right cheek, mouth and chin with different colors, respectively. (B) nose and (C) nose and its adjacent regions example for PWF. (D) angle_EN.N and (E) angle_v example for DAL.

In the second scheme (the phenotypes of distance and angle between landmarks (DAL)), 10 facial features were selected by calculating the distance or angle among 15 landmarks (F1 A). They included 4 nose features (F1 D,E), 4 mouth and chin features, and 2 eye features, details of which were listed in the appendix (method and SF3).

### Genetic association analysis and follow-up study

Genetic association analysis was performed on two different schemes of phenotypes, PWF and DAL (F1), respectively.

For PWF, we performed genome-wide association analysis (GWAS) on the discovery cohort and the replication cohort for the PCs of each facial region. A total of 670273 SNPs was tested in multivariable linear regression analysis by taking gender, age and BMI as covariates under an additive genetic model. We found 42 SNPs among 4 genetic loci that reached the threshold of p < 5 × 10^−8^, which mainly correlated with nose and eyes (ST1). The results of discovery GWAS for each facial region are shown on QQ plots and Manhattan plots with -log10(P value) against the chromosome position (SF4 C-H).

There are 26 SNPs that reached a more conservative threshold 9.32 × 10^−9^ after Bonferroni correction for the facial regions and all genomic loci. Among these eligible SNPs, 17 SNPs were verified in the replication cohort, including 3 SNPs correlated with eyes, 8 SNPs correlated with nose, and 6 SNPs correlated with nose and its adjacent area. All genetic loci were verified in the replication cohort. For further analysis, the SNP with lowest P value in each genetic loci were selected, including rs970797, rs16897517, rs9995821, and rs12637297 (T1). Rs970797, an intergenic SNP near HOXD1 and MTX2 on chromosome 2q31.1(F3 A), was associated with nose (p = 2.15 × 10^−12^) and eyes (p = 1.72 × 10^−8^). Rs16897517, an intronic SNP of VPS13B on chromosome 8q22.2 (F4 B), showed association with nose and its adjacent region (p = 1.39 × 10^−11^). Rs9995821, an intergenic SNP close to DCHS2 and SFRP2 on chromosome 4q31.3 (F4 A), was associated with nose (p = 5.83 × 10^−11^). Rs12636297 near an LncRNA gene PISRT1 on chromosome 3q23 (SF6 A) was associated with eyes (p = 1.73 × 10^−10^).

For DAL, we also conducted GWAS to detect the genetic association of 670273 SNPs with the 10 DAL phenotypes by taking gender, age, and BMI as covariance under the additive genetic model. 5 SNPs (ST2) reached common genome-wide significance (p < 5 × 10^−8^). The results of discovery GWAS for each phenotype are shown on QQ plots and Manhattan plots with -log10(P value) against the chromosome position (SF4 A,B). Only one SNP rs12948076 (p = 7.41 × 10^−9^) around SOX9 and CASC17 gene in the 17q24.3 region met the Bonferroni-corrected significance threshold at 7.45 × 10^−9^ (F2 A), and it was validated for the associations in the replication cohort with angle_v, the angle of Nasion, Pronasale, and Subnasale (landmarks 5, 8, and 9), which represents the length or height of the nose.

**Fig. 2.**
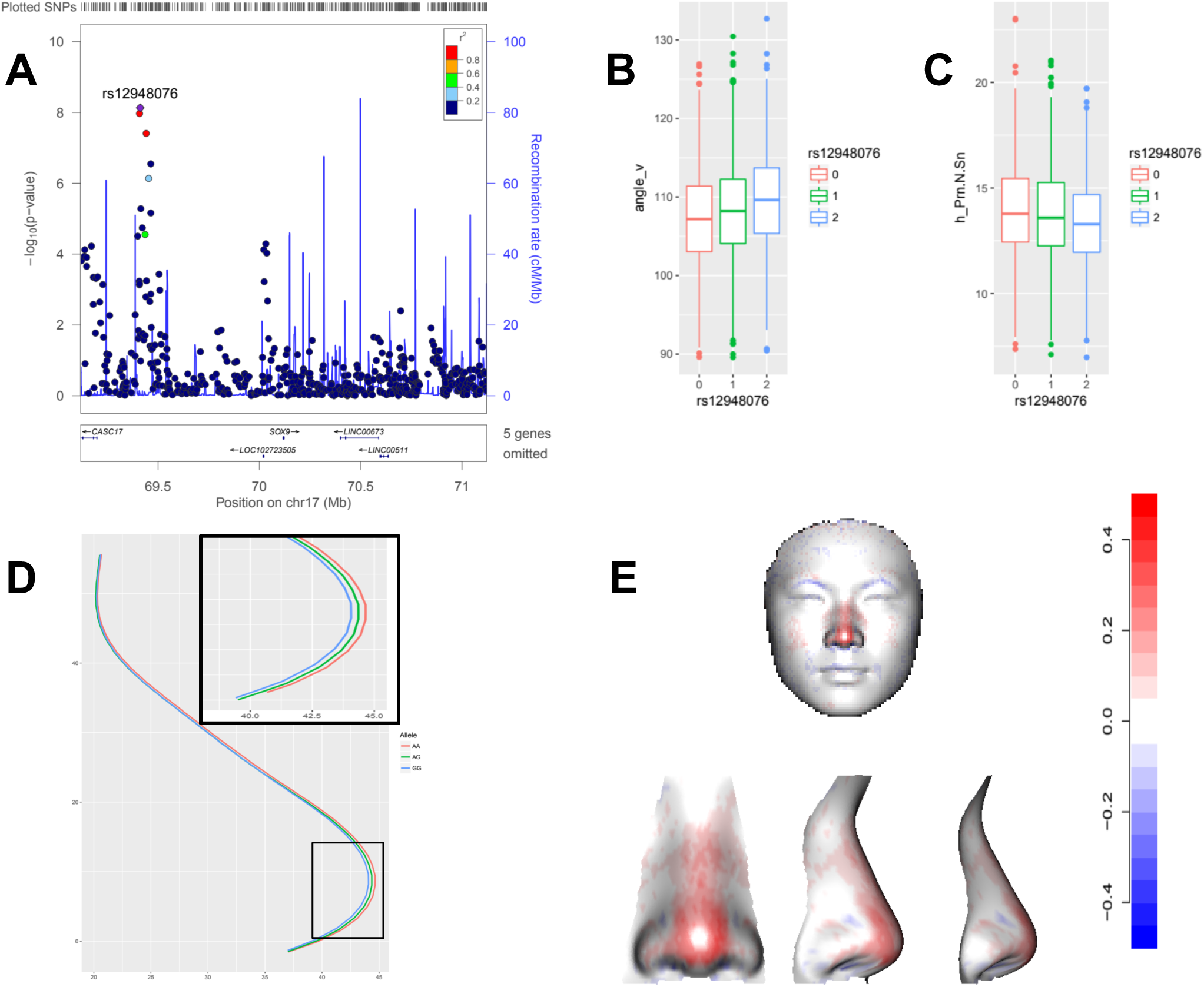
Association between rs12948076 and nasal tip. (A) Locuszoom regional association plot of angle_v at rs12948076 locus in 1 Mb window. Association of SNPs in the discovery GWAS cohort plotted as -log10(P-value) against the chromosomal base-pair position. The y-axis on the left shows -log10(P-value), and the y-axis on the right shows the recombination rate, which was estimated from the HapMap CHB and JPT populations. (B) The boxplot shows the distribution of the angle phenotype angle_v in samples with different genotypes at rs12948076. From left to right, 0 (AA), 1 (AG), 2 (GG) shows in red, green and blue, respectively. (C) the boxplot shows the distribution of the distance phenotype h_Prn.N.Sn in samples with different genotype at rs12948076. (D) the average longitudinal profile of the nose of samples with different genotypes at rs12948076. Zooming into the profile of nasal tip is shown in the solid black wireframe at the upper right. (E) the heatmap of face shows the change of average faces of two homozygous genotypes (AA and GG) at rs12948076 (Details in method), with red indicating outward and blue inward. The following is the heatmap of change in the nose area, which shows in different views, the front, the view of 60 degree, and the side.

**Table 1.**
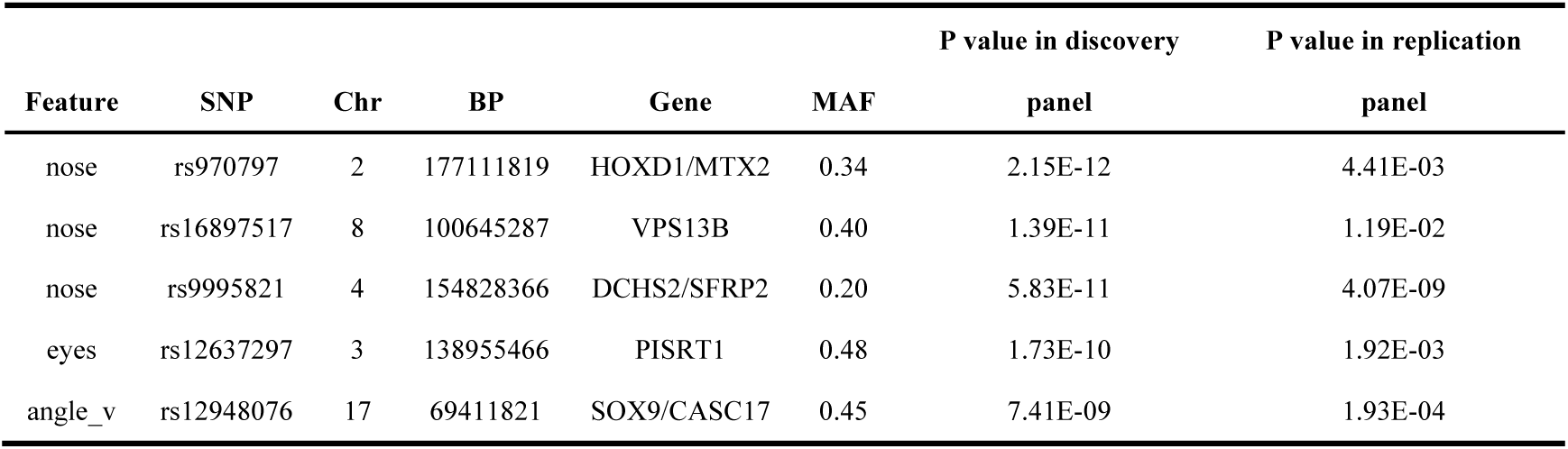
Genome-wide significant loci in discovery cohort and replication cohort.

To further explore the relationship between 4 loci in 17q24.3 with nose, we chose one SNP with the most significant p value (rs12948076,p = 7.41 × 10^−9^) and analyzed its relationship with angle_v. We observed that participants at loci rs12948076 with A allele had smaller angle than with G (F2 B), meaning rs12948076 are related with the length or height of nose and A alleles tend to have shorter nose or higher nose tip than G alleles. Meanwhile, rs12948076 shows some level of association with h_Prn.N.Sn (p = 1.98 × 10^−7^)(ST3), the height of a triangle made up by Nasion, Pronasale, and Subnasale (landmarks 5, 8, and 9), which represents the height of the nose tip. We observed that participants at loci rs12948076 with A allele had higher nose tip than with G (F2 C). In order to intuitively display the effect of different alleles on the nose, we used a novel method to visualize the quantitative changes in the nose area by a 2D axial plane of the nose and a 3D heatmap. From the axial plane of the nose, we noticed that participants at loci rs12948076 with AA had highest nose tip, GG had lowest nose tip, and AG were in the middle (F2 D). Interestingly, there was almost no difference for other parts of the nose. To show the partial and whole effect on nose and face by different alleles, we compared the changes in the volume under each triangle pairs of mean faces, which were separated by rs12948076 under AA and GG alleles. The result showed that participants at rs12948076 loci with AA allele tended to have higher nasal tip than those with GG allele (F2 E). Globally, the genetic variation at rs12948076 may affect the stereoscopic degree of nose, mainly affecting the height of nasal tip, not the other part of face.

We further analyzed the relationship between nose and loci near HOXD1 and MTX2, rs970797. We found that rs970797 has some sort of association with angle_N (p = 4.00 × 10^−7^) and h_N.eye (p = 3.31 × 10^−5^), which are the angle and the height of the triangle made up by right Endocanthion, Nasion, and left Endocanthion (landmarks 2, 5, and 3) (F1 E). These two features represent the height of the nasal root. We observed that participants at loci rs970797 with T allele had larger angle_EN.N (F3 B) and smaller h_N.eye (F3 C) than with G allele, meaning that rs970797 is related to the height of nasion, and T alleles tend to have shorter lower nasion than G alleles. To visualize the difference among TT, TG, and GG alleles, we ploted the 2D axial plane of the nose. It showed that participants at loci rs12948076 with TT had lower nasion than TG and GG alleles, GG alleles had slightly higher nasion than AG alleles (F3 D), consistent with above result. From the 3D heatmap of the changes in the volume under each triangle pair of mean faces separated by rs970797 under TT and GG alleles, we observed that the nasion region was blue (F3 E, SF6), representing the nasion of participants at rs970797 with TT allele more likely to be lower than those with GG allele. Another interesting phenomenon observed from the whole face heatmap is that most regions of the face were sort of blue, indicating that participants at rs970797 with TT tend to have smaller and flatter faces than those with GG allele. In discovery GWAS, we also found that rs97079 met the genome-wide significance level for multiple regions, including nose (p = 8.19 × 10^−8^), eyes (p = 1.72 × 10^−8^), nose with adjacent area (p = 2.15 × 10^−12^) (ST1). The loci rs970797 not only are associated with nasion, but also may affect the stereoscopic degree of the entire face.

**Fig. 3.**
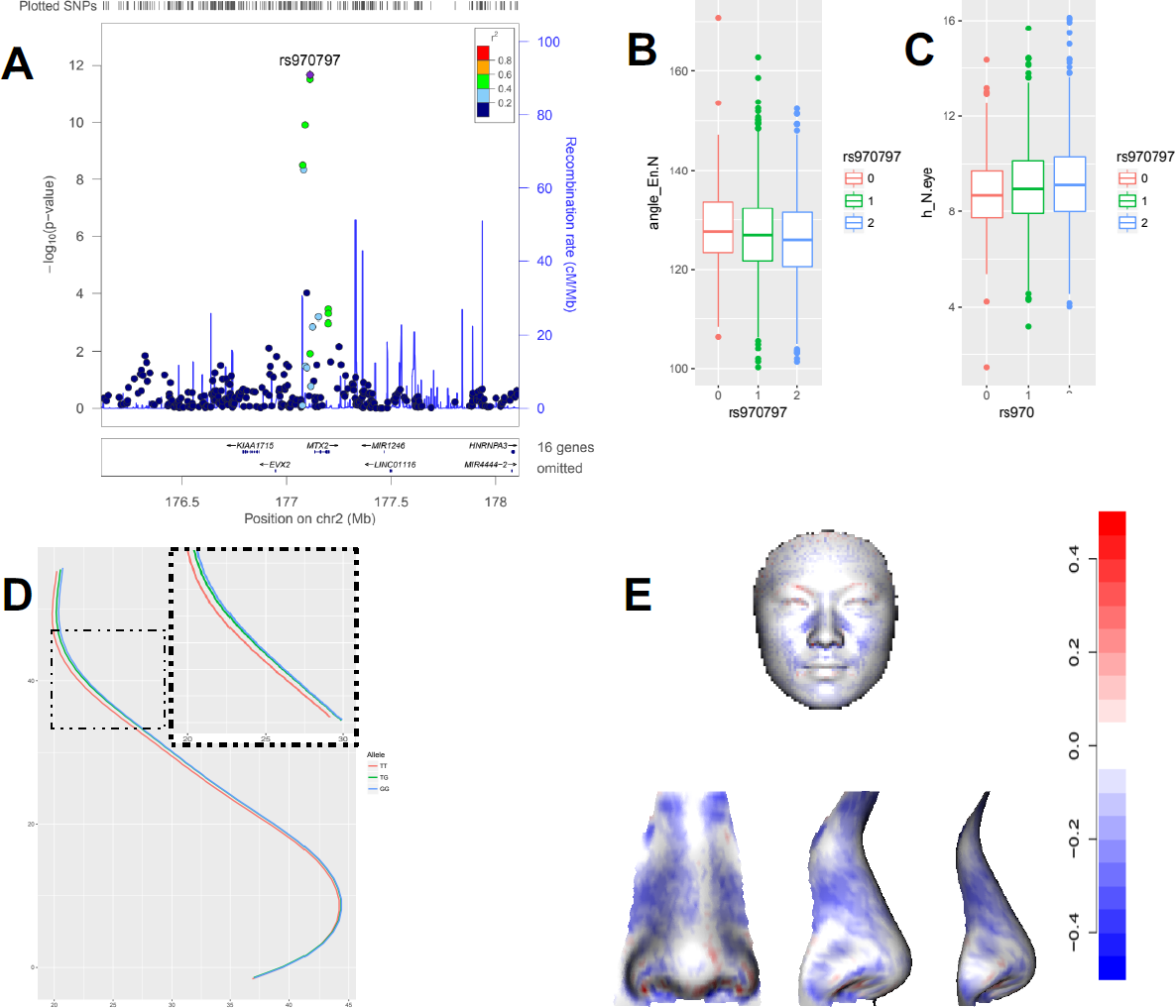
Association between rs970797 and the nasal root. (A) Locuszoom regional association plot of nose and its adjacent regions across a 1 Mb window centered on rs970797. Association of SNPs in the discovery GWAS cohort plotted as -log10(P-value) against the chromosomal base-pair position. The y-axis on the left shows - log10(P-value), and the y-axis on the right shows the recombination rate, which was estimated from the HapMap CHB and JPT populations. (B) The boxplot shows the distribution of the angle phenotype angle_EN.N in samples with different genotypes at rs970797. From left to right, 0 (TT), 1 (TG), 2 (GG) shows in red, green and blue, respectively. (C) the boxplot shows the distribution of the distance phenotype h_N.eye in samples with different genotype at rs970797. (D) the average longitudinal profile of the nose of samples with different genotypes at rs970797. Zooming in the profile of nasal root shows in the dotted black box at the lower left. (E) the heatmap of the face shows the change of average faces of two homozygous genotypes (TT and GG) at rs970797 (Details in method), with red indicating outward and blue inward. The following is the heatmap of change in the nose area, which shows in different views, the front, the view of 60 degree, and the side.

Further analysis revealed that two SNPs rs9995821 and rs16897517 appeared to affect the shape of nostrils (narrowness and length). We compared the mean 3D face of all individuals with CC allele at rs9995821 with the mean 3D face of all individuals with TT allele. Considering the heat map of the differences between the volumes under each pair of triangles (F4 C), we observed that participants at rs9995821 with CC allele tended to have narrower but longer nostrils than those with TT allele. Using the same method, we found that participants with TT allele at rs16897517 tended to have narrower and shorter nostrils than those with GG alleles (F4 D). In the result of discovery GWAS, we noticed that rs16897517 was significantly associated with nose and its adjacent region, while rs9995821 was only significantly associated with nose, indicating that rs16897517 may have a broader influence on the nose region than rs9995821. We also confirmed that rs16897517 had a broader blue region than rs9995821 on the heatmap (F4 C,D). Intriguingly, we observed no difference in the shape of trunk axis curves among different alleles of the two SNPs, and it was difficult to detect the difference, even when amplified (SF5). We think that rs9995821 mainly affects the nostril region, while rs16897517 not only affects the nostril region but also the narrowness of the bridge of nose, but neither of them seems to affect the height of the nose.

**Fig. 4.**
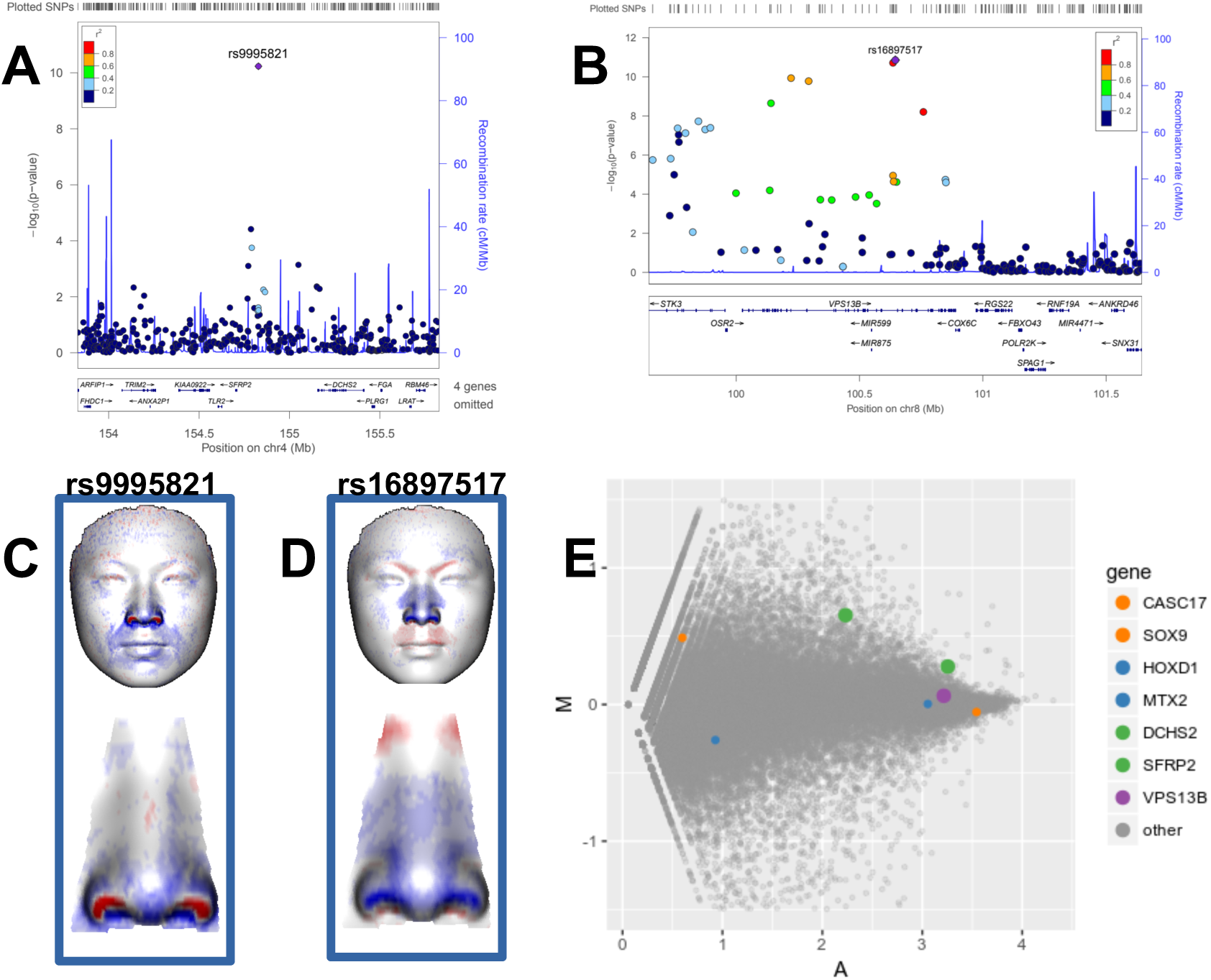
Association between rs9995821 and rs16897517 with the shape of nostrils. (A) Locuszoom regional association plot of nose centered on rs9995821 (B) Locuszoom regional association plot of nose and its adjacent regions centered on rs16897517 (C) At the top, the heatmap of face shows the change of average faces of two homozygous genotypes (CC and TT) at rs9995821, with red indicating outward and blue inward. Zooming-in views are shown in the bottom. (D) At the top, the heatmap of face shows the change of average faces of two homozygous genotypes (TT and GG) at rs16897517, with red indicating outward and blue inward. Zooming-in views are shown in the bottom. (E) The MA plot shows the differences of gene expression in human and chimp CNCCs, by transforming the average gene expression onto M (log ratio) and A (mean) scales. Candidate genes are highlighted.

To explore whether these four nose-related loci and their target genes play an important role in the early development of facial morphology, we compared the epigenetic modifications near the nose-related loci and the expression levels of their target genes by using the data from a public study (Prescott *et al*., 2015) in human and chimp cranial neural crest cells (CNCCs), which plays a crucial role in early facial morphological formation generating individual differences and specially-specific facial features (Bronner and Ledouarin, 2012; Green, Simoes-costa and Bronner, 2015). We observed that there was a strong H3K27ac signal near rs16897517 in the human and chimp CNCCs, and the H3K27ac signal in chimp CNCCs was stronger than in human CNCCs (SF7 B,C). We further analyzed the expression of rs1689517 nearest gene VPS13B in human and chimp CNCCs, and found that VPS13B was highly expressed in CNCC of both species, exceeding the average expression level of 90% genes (F4 E, SF7 A). Although the intensity of enhancer signal of rs9995821 was much lower than that of rs16897517(SF7 B), rs9995821 nearest genes DCHS2 and SFRP2 were both highly expressed, and the expression level of SFRP2 was similar to that of VSP13B (F4 E). Surprisingly, both of DCHS2 and SFRP2 genes tended to be differentially expressed in human and chimp, they had a higher expression in human CNCC (F4 E).

Finally, we further analyzed the relationship between rs12637297 with eyes. From the heatmaps (SF6 B), it seemed that the SNP was mainly associated with the region of eyebrow bone. The result gave an explanation why no significant association with rs12637297 was found in a single eye phenotype.

## Discussion

Recently advanced high-resolution three-dimensional imaging technology enables rapid and accurate acquisition of 3D human facial landscape, which promotes the new possibilities for 3D facial recognition, as well as genetic association studies of human facial morphological variations. Here, a two-stage GWAS of 3D normal human facial morphology in Han Chinese was conducted on two schemes of phenotypes and identified multiple genomic loci significantly associated with nose-related traits and eye-related traits. Through analyzing the relationship between phenotypes and genotypes and visualizing the changes on facial morphology, we further refined the effects of these genetic loci on different parts of the nose.

Salience of nasal tip: rs12948097 with A allele likely protrudes the nasal tip. The SNP tends to affect nose prominence, but not the whole face. There are two genes, SOX9 and CASC17, that are close to rs12948076. It is very interesting that SOX9 and CASC17 as two candidate genes at 17q24.3 have also been reported in the latest GWAS of facial morphology with a data-driven method to phenotyping facial shape (Claes *et al*., 2018). SOX9 was associated with the prominence of the nose tip, and CASC17 was also associated with the prominence of the nose tip, along with the sidewall of the nose. Another GWAS of normal facial morphology based on 2D photographs of Korean individuals reported the association between SOX9 with multiple nose-related phenotypes, such as nasal tip protrusion, and nasal bridge depth (Cha *et al*., 2018). We observed that SOX9 had very high expression in human and chimp CNCCs, but CASC17 had very low expression. This indicates that SOX9 is more likely to be the target gene of rs12948076. Previous studies described a key role of the SOX9 gene in chondrogenesis considered as the “main regulator”, and showed that the combination of SOX9 with other transcription factors can activate the transcription of the chondrocyte-specific factor COL2A1 and promotes the differentiation of mesenchymal stem cells into chondrocytes (Lefebvre and de Crombrugghe, 1998; Wilde, 2009; Ahmed *et al*., 2014). A mutation of the SOX9 gene can lead to a rare disease of bone dysplasia, campomelic dysplasia (CMD), and most of the patients with CMD have dysplasia of the facial endochondral bone, resulting in a variety of facial deformities, such as cleft palate, depressed nasal bridge (Tommerup *et al*., 1993; Mansour *et al*., 2002). In addition, the SOX9 gene is involved in normal osteoblast differentiation and bone formation by limiting RUNX2 expression, and a SOX9 mutation could cause cleft palate due to premature palatal development (Gopakumar *et al*., 2014; Yao *et al*., 2015).

Size of nasal root: rs970797 with T allele tends to flatten the nasal root. Two recent GWAS of normal human face found that the exact same locus rs970797 was associated with lip prominence (Claes *et al*., 2018) and eye shape (Cha *et al*., 2018), respectively. These confusing results were intuitively explained in our study. We found that rs970797 may affect the stereoscopic degree of the whole face from the global perspective. Participants with TT allele at rs970797 tend to have smaller and flatter face than those with GG allele. HOXD1 and MTX2 are two candidate genes near rs970797. HOXD1 did not express in human and chimp CNCCs (F4 E). Reviews have summarized that the developing face does not express the HOX genes which widely express in other regions to determine anterior-posterior patterns (Mallo, Wellik and Deschamps, 2010; Casaca, Santos and Mallo, 2014). MTX2 becomes an interesting candidate gene due to the lacking expression of HOXD1 in cranial neural crest cells.

Shape of nostrils: rs16897517 with T allele is associated with shorter nostrils, and rs9995821 with C allele with longer nostrils. These two SNPs seem to primarily influence the shape of nostrils. DCHS2 and SFRP2 are the candidate genes of rs9995821, and VPS13B is the candidate gene of rs16897517. the SNP rs9995821 had been reported to be associated with ala aperture and nostrils (Claes *et al*., 2018), which was identical with our results. Meanwhile, multiple loci near DCHS2 were associated with columella inclination and nose tip angle in admixed Latin Americans (Adhikari *et al*., 2016). DCHS2 was first discovered in Drosophila and plays an important role in tissue morphogenesis and growth control(Clark *et al*., 1995; Zecca and Struhl, 2010). A recent study found that it is involved in the regulation network of cartilage differentiation during vertebrate craniofacial development (Le Pabic, Ng and Schilling, 2014), which is initiated by SOX9 (Bi *et al*., 1999). Another candidate gene SFRP2 is a negative regulator of WNT signaling (Kurosaka *et al*., 2014). Disrupting hedgehog and WNT signaling interactions promotes cleft lip pathogenesis, and craniofacial deformities have been reported in SFRP2 mutant mice (Kurosaka *et al*., 2014). Double homozygous mutations in Sfrp1 and Sfrp2 result in severe shortening of the thoracic region (Satoh, 2006). Remarkably, DCHS2 and SFRP2 were differentially expressed in human and chimp CNCCs, which might make the human nose big and thereby unique among primates (Nishimura *et al*., 2016). We found the new locus, rs16897517, located in the intron of VPS13B. VPS13B was also found in the discovery cohort of the Korean population, although it was not successfully verified in the replication cohort (Cha *et al*., 2018). It is in a very strong signal enhancer region in both human and ape CNCCs, and its candidate gene VPS13B were highly expressed in both species. Mutations in VPS13B gene underlie Cohen syndrome, a rare autosomal recessive disorder with characteristic craniofacial dysmorphism, such as micrognathia, short philtrum and high vaulted palate (Balikova *et al*., 2009; Seifert *et al*., 2009). Through the analysis of epigenomic profiling and gene expression of human and chimp CNCCs, we found two SNPs, rs16897517 with a strong enhancer signal in VPS13B and rs9995821 with differential expression of candidate genes, may be related to differences of the nose in human and chimp.

In conclusion, we defined two groups of phenotypes to characterize facial morphology in 2659 Han Chinese and found four nose-related SNPs as well as one eye-related SNP (SF8). Three of the seven candidate genes and two loci were reported in the latest GWAS of facial morphology with a data-driven method to phenotyping facial shape. One candidate gene was reported in admixed Latin Americans. Furthermore, four candidate genes were independently found in a recent GWAS of normal human facial morphology in the Korean population, and two of these related loci were identical. Populations with similar genetic backgrounds are more likely to find overlapping results than those with different genetic backgrounds, and the data-driven segmentation method has the advantage of dealing with human face. We visualized the effect of genetic variants on different parts of the nose, which resolved the fuzzy results of different phenotypes and showed information beyond the phenotype. We noted that the nose underlies precise regulation of genetic variants. These results provide a more holistic understanding of the relationship between genetic loci with human facial morphology, which potentially play a role in forensic face prediction, facial disease prediction, and evolutionary implications.

## Materials and Methods

### Sample Information

The 3,054 samples used in this study were collected in Taizhou, Jiangsu province in 2014, among which 1,091 were males and 1963 were females. The three-dimensional facial images of the participants were collected by a 3dMDface 3d camera system, and blood samples from participants were also collected. In addition, the age, height and other information of the sample were collected through questionnaires. Samples were aged 31 through 87 (mean: 56.7, sd: 9.5), with males’ age 34-87 years old (mean: 58, sd: 9.8) and females’ age ranging between 31-86 years old (mean: 56, sd: 9.3) (detail supfig s1). The BMI of samples were ranging between 14.7 and 50.3 (mean: 24.4, sd: 3.2), with males’ BMI ranging between 14.7-35.2 (mean: 24.6, sd: 3) and females’ BMI ranging between 15.6-50.3 (mean:24.3, sd: 3.3) (detail supfig s2). All participants were apparently unrelated Han Chinese residents, with no history of facial trauma, facial plastic surgery, or any known disease affecting the face. In addition, participants were excluded if they had any personal or family history of facial deformities or birth defects. After removing 395 participants, with missing personal information, 3D image mapping artifacts, or unqualified control of GWAS genotyping, a total of 2659 participants were retained for analysis.

### Ethics statement and consent to participate

All participants provided written informed consent to participate in the project. Sample collection for this study was conducted with the approval of the ethics committee of Shanghai Institutes for Biological Sciences and in accordance with the standards of the Helsinki Declaration.

### High-density 3D facial image collection and registration

The workflow of overall procedure is shown in supfig 3. There are 4 steps: capturing 3D image by 3dMDface camera system, landmarking critical points of the face by Facial Registration Analysis Software (FRAS), mapping with reference facial mesh, and adjusting coordinate system by Generalized Procrustes Analysis (GPA).

Capturing 3D image by 3dMDface camera system. 3D facial image of participants in this study were all captured by the 3dMDface system (www.3dmd.com/3dMDface), which captures a 3D facial image at a speed of ∼ 1.5 milliseconds with a geometric accuracy of 0.2 mm RMS (Weinberg *et al*., 2016). According to standard facial image acquisition protocols (Heike *et al*., 2010), participants were required to follow the following conditions: having a natural facial expression without wearing any cosmetic make-up, pulling back the hair covering the forehead, opening eyes without wearing glasses, and closing the mouth naturally. After the shooting process, the quality of the 3D images was double checked on the computer through the software provided by 3dMDface. 3D images were saved in TSB format by the 3dMDface system and converted to OGJ format for later processing.

Landmarking critical points of the face by Facial Registration Analysis Software (FRAS). 17 critical points were automatically annotated on the meshes using an in-house 3D face landmarking algorithm (FRAS) based on PCA projection of texture and shape information to achieve the high-density point-to-point registration (Guo, Mei and Tang, 2013). In briefly, the OBJ files were imported into Facial Registration Analysis Software. The nose tip was first automatically identified by using a sphere fitting approach. Then, the pose was normalized into a uniform frontal view. The postural standardized 3D facial images were projected into two-dimensional space through PCA, and the remaining 16 points are annotated automatically according to the geometric relationship and texture information (Guo, Mei and Tang, 2013). These landmarks were checked in FRAS and corrected by 3dMD Patient if the landmark was an outlier.

In this study, 2 landmarks at two earlobes were removed because many female participants blocked their ears by loose hair.

Mapping with reference facial mesh. A facial mesh with high-image quality and smooth surface was selected from the annotated faces and resampled uniformly to achieve an even density of one vertex per 1mm-by-1mm grid area. The reference face was a high-density grid mesh with 32251 vertices. Then the tool Thin Plate Spline (TPS) was used to twist and distort each face in the sample in turn, and the 15 landmarks on each surface were aligned with those on the reference face in a non-rigid way. Each vertex in the reference face was mapped to the closest point on the sample faces, and the sample faces form a high-density mesh that matches the reference face. Each surface is represented by 32251 points of 3D mesh.

Adjusting the coordinate system by Generalized Procrustes Analysis (GPA).

In order to eliminate differences in position and orientation of surfaces, all surfaces of participants were shifted and rotated to the same coordinate system by Generalized Procrustes Analysis (GPA).

During the processing of the above steps, the sample images were screened to remove the samples that had a serious defect in the facial images and did not pass the above steps due to other reasons. Finally, the remaining 2,963 samples with high-quality images obtained the coordinate information in the same coordinate system through the above workflow for subsequent GWAS analysis. Each face is represented by a tri-mesh through a point matrix of dimensions 32251 by 3 and the edge information of these points.

### Phenotyping quantitative features from the tri-mesh facial surface

Two schemes of quantitative phenotype were used to describe facial morphological features in this study.

The first scheme (the partial and whole facial phenotype) used geometric analysis of high-density data. According to the anatomy of the face, the facial surface was divided into the areas of eyes, cheeks, nose, mouth and jaw, and the partial face of the corresponding areas were extracted from the high-density tri-mesh surface by FARS and 3dMD Patient software. Marker points of each partial face were first superimposed into a common coordinate system by GPA. Then PCA was performed on the high-density tri-mesh surface of partial face, and 97 percent of PCs with the total variance of interpretation were extracted to characterize the morphological features of each region. In addition, PCA was also performed on the whole face to obtain PCs explaining 97 percent of the total variance, to represent the changes of facial morphology from the overall perspective. Selected PCs were taken as phenotypes in Multivariable Linear Regression (MLR).

Another scheme (the distance and angle phenotype) was based on 15 landmarks obtained from the high-resolution tri-mesh facial surface. 10 facial traits composed by 15 landmarks were provided in Supplementary Figure 3. For example, 6-8-7 means the distance between Landmarkss 6 and 8 plus the distance between Landmarks 8 and 7. Angle 6-8-7 means the angle between the line through Landmarks 6 and 8, and the line through Landmarks 7 and 8.

### Genotyping, quality control, and population structure

Participants including 2978 Han Chinese from Taizhou, Jiangsu Province and 2 Uyghur from Urumchi, XinJiang Province were genotyped on Illumina Omni ZhouHua-8, and a total of 887,270 SNPs sites were obtained. Quality control filtering of genome-wide genotype data was performed by using PLINK v1.07 (Purcell *et al*., 2007).

The quality control process was mainly divided into two steps.

1. Quality control of chip data. SNP loci were required to satisfy the following conditions:
  A. The Marker Missing rate is less than 0.02.
  B. Minor Allele Frequency (MAF) is greater than 0.05.
  C. Hardy-Weinberg Equilibrium test with a p-value of less than 0.001.
  D. Exclusion of SNPs without chromosome information and duplicated SNPs. After removing the unqualified SNPs, there were 670273 remaining SNPs.
2. Quality control of samples. The samples were required to meet the following conditions:
  A. Individual missingness. 4 participants were removed due to low calling rate after SNPs pruning.
  B. Duplicates and cryptic relatedness. 2 pairs of participants had extremely high IBD, which means that the two participants of each pair may be identical.
  C. Population structure. 2 Uyghur participants were far away from the center of Han participants.
  D. Heterozygosity and inbreeding. None of the participants were removed at threshold value > 0.2 or < - 0.2)
  E. Gender check. For 4 individuals, the gender did not fit the information given (F value less than 0.2 for female, more than 0.8 for male).

### Statistical Analysis

GWAS was performed on 2959 participants with 670273 SNPs (952 male and 1707 female) and two schemes of phenotype (the distance and angle phenotype, and the partial and whole face phenotype).

Multiple linear regression analysis was performed to discover variants associated with each of the distance and angle phenotype, under an additive genetic model, while simultaneously adjusting for age, BMI and gender. For the partial and whole face phenotype, multivariable linear regression analysis was performed for genetic association between each SNP with PCs of nose, eyes, and mouth. The prcomp function in the R statistics package was used for PCA to extract the variance on the high-density tri-mesh surface.

Similar analyses were also applied to the association tests of the indexed SNPs in the replicated cohorts.

Quantile-quantile plots for facial phenotypes were shown with the distribution of observed P-values against the theoretical distribution of expected P-values through qqman.

The manhattan plots for facial phenotypes with significant association loci were displayed with -log10(P-value), the SNPs with significant signal were highlighted, and two threshold lines at 5 × 10^−7^ (Orange) and 5 × 10^−8^ (Red) were drawn by CMplot. The regional association plot for a genomic region of 1 Mb centered on the significant loci was constructed using LocusZoom.

There were two kinds of mathematical models for genome-wide association analysis (GWAS) in this study.

(1) Multiple Linear Regression

Y_phenotype ∼ X_snp + X_age + X_bmi + X_gender

Y_phenotype represented the distance or Angle between landmarks.

X_snp refered to the genotype of a particular SNP in the sample (0, 1, 2).

X_age was the age information of the sample.

X_bmi was the BMI information of the sample.

X_gender was the gender information in the sample, 1 for male and 2 for female.

The quantitative features of each facial morphology described by distance or angle were correlated with all the SNPs. The additive model based on multiple linear regression was implemented by encoding genotypes as 0, 1, and 2. T test was used to determine whether an SNP had a significant effect on facial morphology with a strict significance threshold of P<5×10-8, given the burden of multiple comparisons.

(2) Multivariable Linear Regression

Y_pc1+Y_pc2+ … + Y_pcm ∼ X_snp+X_age+X_bmi+X_gender

Y_pc1, Y_pc2 and Y_pcm were the PCs extracted from the partial or whole facial phenotype.

X_snp, X_age, X_bmi and X_gender were the same as described above in (1).

For a certain region of the facial division, the corresponding PCs and all the SNPs were analyzed one by one according to the above model. The significance of an SNP was tested by the Pillai-Bartlett test.

### Face quantification and visualization

Each 3D human face was represented by a tri-mesh surface with 32251 points after elastic registration to the reference face. the tri-mesh contains 63770 triangles. all surfaces were under the same coordinate system by Generalized Procrustes Analysis (GPA). Then, the average faces of the genetic loci of interest were calculated according to different genotypes, such as TT, TG, and GG, and the volumes of tetrahedrons formed by the origin and triangle in the average surfaces were calculated, respectively. Since GWAS was based on an additive model in the study, it is considered that these genetic variants also have an additive effect on the phenotype. The changes of the local face were obtained by correspondingly subtracting the volumes of the average face of homozygous genotypes and dividing the mean of distance to origin as normalization. To quantify and visualize the effect of genetic variants on the human face, we developed an open source R package “VFave” based on rgl. VFace provides functions to read a 3D face object, calculate the volume of tetrahedrons formed by the origin and a triangle in the tri-mesh surface, and draw a 3D face with different style and gradient colors. Source code can be accessed in GitHub (https://github.com/changebio/VFace).

### The analysis of epigenomic profiles and gene expression in human and chimp CNCCs

Epigenomic and expression data were collected from Prescott et al., including histone modifications (H3K4me1, H3K4me3, H3K27ac,and H3K27me3) ChIp-seq, transcription factors (TFAP2A and NR2F1) ChIp-seq, a general coactivator (p300) ChIP-seq, and chromatin accessibility ATAC-seq, as well as gene expression RNA-seq from both human and chimp CNCCs. We focused on cis-regulatory activities near associated loci and gene expression of candidate genes. Normalized H3K27ac coverage was calculated in the 20-kb genomic regions centered on the 5 associated genetic loci, and a box plot of the normalized H3K27ac signal with multiple replicates in human and chimp CNCCs was plotted using ggplot2. The gene expression was normalized by variance stabilizing transformation (VST). The box plot of gene expression of candidate genes with multiple replicates in human and chimp CNCCs was plotted using ggplot2. The MA-plot visualized the differences between average gene expression in human and chimp CNCCs by transforming the average expression of genes onto M (log ratio) and A (mean) scales, with highlighted candidate genes. The signal track of histone modification and TFs was shown in the 20-kb genome region centered by the associated SNP by uploading wig files to the UCSC genome browser (Kent *et al*., 2002).

## Supplementary Information

Supplementary information includes figures and tables.

## Acknowledgements

The authors are grateful to all the volunteers taking part in this study. We would like to thank Yu Liu, Qianqian Peng, Sijie Wu, and Manfei Zhang for their role in data collection and genomic data cleaning; thank Lu Qiao for the image processing. This work was funded by the Max-Planck-Gesellschaft Partner Group Grant (KT), the National Natural Science Foundation of China (Nos. 31371267, 31322030, 91331108 (KT); 91631307 (SW); 30890034, 31271338 (LJ); and 91731303, 31771388, 31711530221 (SX)). This work was supported by the National Basic Research Program (2015FY111700, LJ), Shanghai Municipal Science and Technology Major Project (2017SHZDZX01, LJ), the Ministry of Education (311016, LJ), Strategic Priority Research Program of the Chinese Academy of Sciences (CAS) (XDB13040100, SX; XDB13041000, SW), the National Science Fund for Distinguished Young Scholars (31525014, SX), the Program of Shanghai Academic Research Leader (16XD1404700, SX), the support of a National Thousand Young Talents Award and a Max Planck-CAS Paul Gerson Unna Independent Research Group Leadership Award (SW), the Science and Technology Commission of Shanghai Municipality (16JC1400504, SW).

## Author Contributions

Conceived and designed the study: K.T., S.G. Performed the study, summarized results and wrote the manuscript: Y.H., D.L. Data processing: L.Q., Y.L., Q.P., S.W., M.Z. Advise the study and modified the manuscript: K.T., S.G., S.W.

